# Dynamic ASK1 proximity networks uncover SCF-dependent and noncanonical roles in ABA and drought adaptation

**DOI:** 10.64898/2025.12.22.696057

**Authors:** F. Daniela Rodriguez-Zaccaro, Jacob Moe-Lange, Shikha Malik, Christian Montes, Natalie Hamada, Andrew T. Groover, Justin W. Walley, Nitzan Shabek

**Affiliations:** Department of Plant Biology, College of Biological Sciences, University of California-Davis, CA; Department of Plant Pathology, Entomology and Microbiology, Iowa State University, Ames, Iowa; Northern Research Station, USDA Forest Service, Burlington, Vermont

**Keywords:** Ubiquitin-Proteasome System, Abscisic Acid (ABA), Drought Adaptation, Proximity Labeling, Quantitative Proteomics, Plant adaptive signaling

## Abstract

Plants rely on rapid proteome remodeling to withstand fluctuating environmental conditions, yet how the ubiquitin system dynamically coordinates these multilayered responses remains unclear. Here we define the in vivo proximity interactome of ARABIDOPSIS SKP1-LIKE 1 (ASK1), the core adaptor of SCF ubiquitin ligases, under acute abscisic acid (ABA) signaling and prolonged drought. TurboID-based proximity labeling coupled with quantitative proteomics revealed that ASK1 assembles highly condition-specific protein networks, distinguishing canonical SCF modules from broader noncanonical associations with transcriptional, chromatin, translational, vesicle-trafficking, and proteostasis machinery. Acute ABA exposure rapidly recruits F-box proteins and ABA-responsive transcription factors while engaging ribosomal and chromatin modules, whereas drought drives ASK1 into expanded proteostasis and stress-signaling assemblies, including chaperone-cochaperone systems, transcriptional repressors, and autophagy-endomembrane components. Global proteomics shows that ASK1 overexpression enhances accumulation of drought-protective and ABA-responsive proteins while repressing immune and ROS-scavenging pathways, indicating a shift in resource allocation. Together, these results describe ASK1 as a multifunctional proteostasis and signaling hub that integrates SCF-dependent and SCF-independent pathways to coordinate transcriptional, translational, and proteolytic reprogramming during plant adaptation to stress.

## Introduction

Plants have evolved elaborate signaling networks to coordinate growth and development with fluctuating environmental conditions. To survive drought, salinity, and other stresses, plants must rapidly adjust metabolic and developmental programs, a process that depends on dynamic proteomic plasticity. This plasticity requires tight, temporally controlled turnover of key regulators such as receptors, transcription factors, and biosynthetic enzymes, largely mediated by the ubiquitin-proteasome system (UPS) (Vierstra 2009). Plant genomes encode a vast number of ubiquitin E3 ligases, with over 1000 reported in *Arabidopsis thaliana* alone, significantly surpassing the numbers found in animal and yeast systems (Mazzucotelli et al. 2006). Among the diverse families of E3 ligases, Cullin-RING Ligases (CRLs) form the largest and most versatile superfamily. As multi-subunit complexes, CRLs are modular multi-subunit complexes built around a cullin scaffold that recruits an E2 ubiquitin-conjugating enzyme via the RING protein RBX1 at its C terminus, and a substrate-recognition module at its N terminus composed of adaptor (e.g., SKP1 or BTB) and receptor (e.g., F-box or DDB2) proteins (Zheng and Shabek 2017; Shabek and Zheng 2014). The SCF (SKP1-CUL1-F-box) represents the archetypal CRL and a paradigm for understanding E3 ligase modularity. The extraordinary expansion of F-box genes in plants (∼700 in Arabidopsis) provides a vast repertoire of substrate receptors, enabling SCF complexes to integrate developmental, hormonal, and environmental cues (Kelley 2018; Tal et al. 2022; Tal et al. 2020). ABA is a key phytohormone governing adaptive processes such as stomatal closure, osmolyte accumulation, and transcriptional activation of stress-responsive genes (Kavi Kishor et al. 2022). Mounting evidence from plant genetics and phenotypic analyses identify SCF complexes as pivotal in fine-tuning abscisic acid (ABA) signaling in response to drought and salinity (Cheng et al. 2017; Li et al. 2012).

Within the SCF complex, ASK1 (ARABIDOPSIS SKP1-like) adaptor proteins play an indispensable role by linking F-box receptors to CUL1 scaffold. Unlike mammals which typically have one or two copies of SKP1-like genes, plants have experienced extensive gene duplication with the Arabidopsis genome encoding 21 ASK proteins (Farras et al. 2001). Among these, ASK1 is the best characterized and its functional importance is underscored by the severe developmental deficiencies, including male meiotic defects observed in *ask1* knockout mutants (Yapa et al. 2020; Zhao et al. 2003; Yang et al. 1999). Similarly, mouse SKP1 localizes to synapsed chromosomes and is necessary for normal meiosis (Guan et al. 2020). Higher order mutants are embryo lethal, making it challenging to study *ASK* roles in stress responses through traditional genetic loss-of-function approaches (Yapa et al. 2020). Knockdown of ASK1 in Arabidopsis confers ABA hyposensitivity, as evidenced through germination and root growth assays (Li et al., 2012; Yappa et al., 2020). Intriguingly, recent work in mammals revealed that SKP1 can also exert noncanonical functions independent of the SCF complex, acting as a molecular switch between autophagy and unconventional secretion under nutrient stress (Li et al. 2023). Such stress conditions impose contrasting proteomic demands: acute ABA exposure triggers rapid signaling reconfiguration, whereas prolonged drought demands sustained proteome remodeling to maintain cellular homeostasis. E3 ligases, particularly SCF complexes, are prime mediators of such plasticity because they can swiftly alter the stability of signaling components. The dual potential of ASK1, functioning both within the SCF and independently of it, makes it a compelling candidate for coordinating canonical and alternative proteome responses across short- and long-term stress conditions (Fig. 1A).

**Figure 1.**
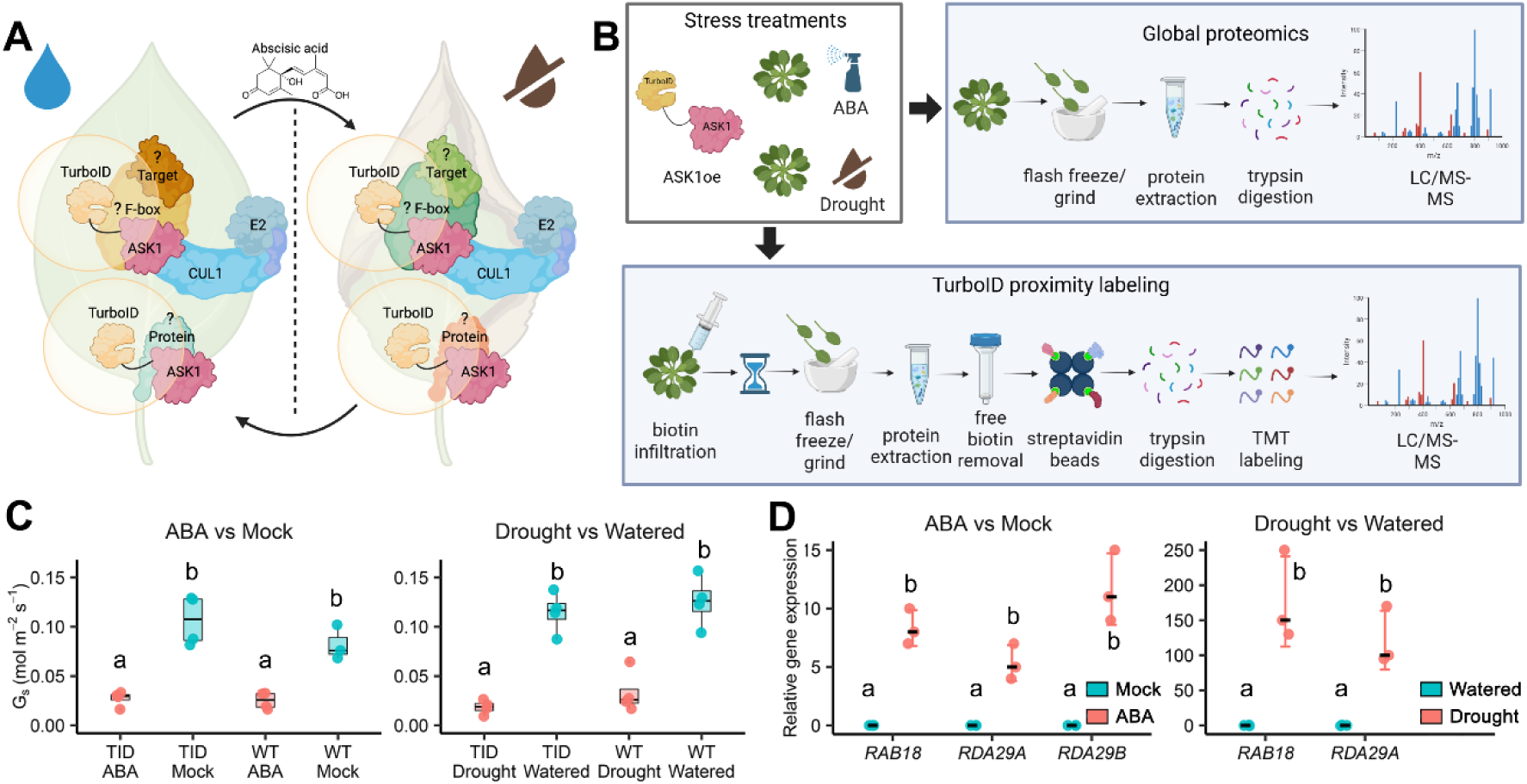
Overview of abscisic acid (ABA) and drought treatments in TurboID-ASK1 Arabidopsis plants for proteomic, physiological, and gene expression analyses. **A)** Schematic of our working model proposing ABA/drought-dependent shifts in ASK1-proximal interactions. Under mesic conditions, TurboID-ASK1 may label nearby SCF components, F-box proteins, putative substrates, or ASK1-associated proteins. With increased ABA, we hypothesize changes in the recruitment of F-box proteins and candidate substrates, altering the ASK1-proximal proteome. **B)** An ABA (50 μM spray) or drought treatment (20% of field capacity) was applied to Arabidopsis plants expressing TurboID-ASK1. Separate groups of treated individuals were collected for either global proteomic analysis to obtain protein abundance profiles or TurboID proximity labeling. **C)** Leaf stomatal conductance (Gs) was measured in ABA-treated versus mock controls and in drought-treated versus well-watered TurboID-ASK1 and wild-type (Col-0) plants. **D)** The expression of ABA marker genes in both ABA and drought-treated TurboID-ASK1 plants was investigated through rt-qPCR analysis. Groups that share a letter are not significantly different (Adj.P Tukey >0.05). Boxes indicate the interquartile range (IQR), the center line denotes the median, and whiskers represent the data range. Parts of this figure were created with BioRender.com.

Given that ASK1 is a necessary component for core developmental processes, stress signaling, and may harbor functions independent of the canonical SCF complex, a comprehensive understanding of its protein interaction landscape is crucial. To determine whether ASK1 engages in distinct molecular functions during both acute (short-term) and chronic (long-term) drought stress, we applied TurboID-based proximity labeling, an in vivo proteomic approach that captures transient and stable protein-protein associations (Sun et al. 2024; Cho et al. 2020). By integrating these proximity networks with global protein abundance profiling, we uncovered extensive stress-dependent remodeling of ASK1-centered assemblies. The resulting datasets revealed previously unreported ASK1-proximal proteins and pathways and illustrate that ASK1’s interaction landscape is dynamically remodeled under different stress regimes. The combined analyses indicate that ASK1 is positioned within multiple stress-responsive protein networks and highlight its potential to coordinate broad proteome remodeling during plant acclimation to environmental stress.

## Results

### ABA and drought treatments establish a defined physiological context for ASK1 network analysis

To investigate the dynamic role of ASK1 complexes in abiotic stress adaptation (Fig. 1A), we employed TurboID-based proximity labeling to reveal its in-vivo interaction networks under short-term ABA signaling and prolonged drought stress. Transgenic Arabidopsis thaliana lines expressing UBQpro:TurboID-3xMyc-gASK1, previously validated for efficient in planta biotinylation of ASK1-associated proteins (Sun et al. 2024) were used for both proximity labeling and global proteomic analyses. Plants were treated with either a 50 µM ABA spray, with tissue harvested three hours later, or exposed to drought stress by allowing soil to dry to 20% field capacity over approximately three weeks, after which shoot tissues were collected (Fig. 1B). To confirm that our ABA and drought treatments elicited robust stress responses, we monitored leaf stomatal conductance (Gs) and examined the expression of canonical ABA-responsive marker genes. Both TurboID-ASK1 and wild-type (Col-0) plants showed significantly reduced stomatal conductance two hours after ABA treatment compared to mock-treated controls (Fig. 1C). Similarly, prolonged drought stress led to significantly lower leaf stomatal conductance values relative to well-watered plants across both genotypes (Fig. 1C). These effects were independent of genetic background, indicating that the TurboID-ASK1 transgene did not alter baseline or stress-induced stomatal behavior. At the molecular level, ABA treatment significantly induced the expression of RAB18, RDA29A, and RDA29B, well-established markers of ABA signaling compared to mock treatment (Fig. 1D). Under drought conditions, RAB18 and RDA29A also showed strong upregulation relative to well-watered controls, (Fig. 1D), consistent with sustained ABA pathway activation. Together, these physiological and molecular responses confirm that both acute ABA exposure and prolonged drought effectively triggered stress signaling in the system, providing a reliable framework for subsequent proteomic analyses.

### ABA signaling triggers rapid remodeling of the ASK1 proxitome

Following validation of stress activation, we next examined how ASK1 complexes reorganize during acute ABA signaling using TurboID-based proximity labeling coupled with quantitative TMT proteomics (Fig. 1B). Across mock and ABA treatments, we identified 1,465 proteins in the ASK1 proxitome, with 425 proteins exhibiting significant differential enrichment (FDR < 0.05 and |log_₂_FC| ≥ 0.27). Because TMT ratio compression tends to underestimate fold changes, this cutoff likely reflects more substantial differences in vivo (Ting et al. 2011; Hogrebe et al. 2018). Among the differentially abundant proteins, 276 were enriched under ABA treatment and 149 under mock conditions, suggesting a marked ABA-dependent expansion and compositional shift of ASK1-associated complexes (Fig. 2A). Functional enrichment of Gene Ontology (GO) biological process categories revealed distinct ASK1 interaction profiles under mock and ABA treatments. Under mock conditions, the ASK1 proxitome was enriched for 13 GO biological processes, reduced to four non-redundant categories including jasmonic acid (JA) signaling and negative regulation of defense responses (Fig. 2B). These included the JASMONATE-INDUCED OXYGENASES 2, 3, and 4 (JOX2, JOX3, and JOX4) which are known to inactivate jasmonic acid via hydroxylation (Caarls et al. 2017). Protein complex oligomerization, localization within membrane and response to light intensity-related annotations were also significantly enriched (Fig. 2B).

**Figure 2.**
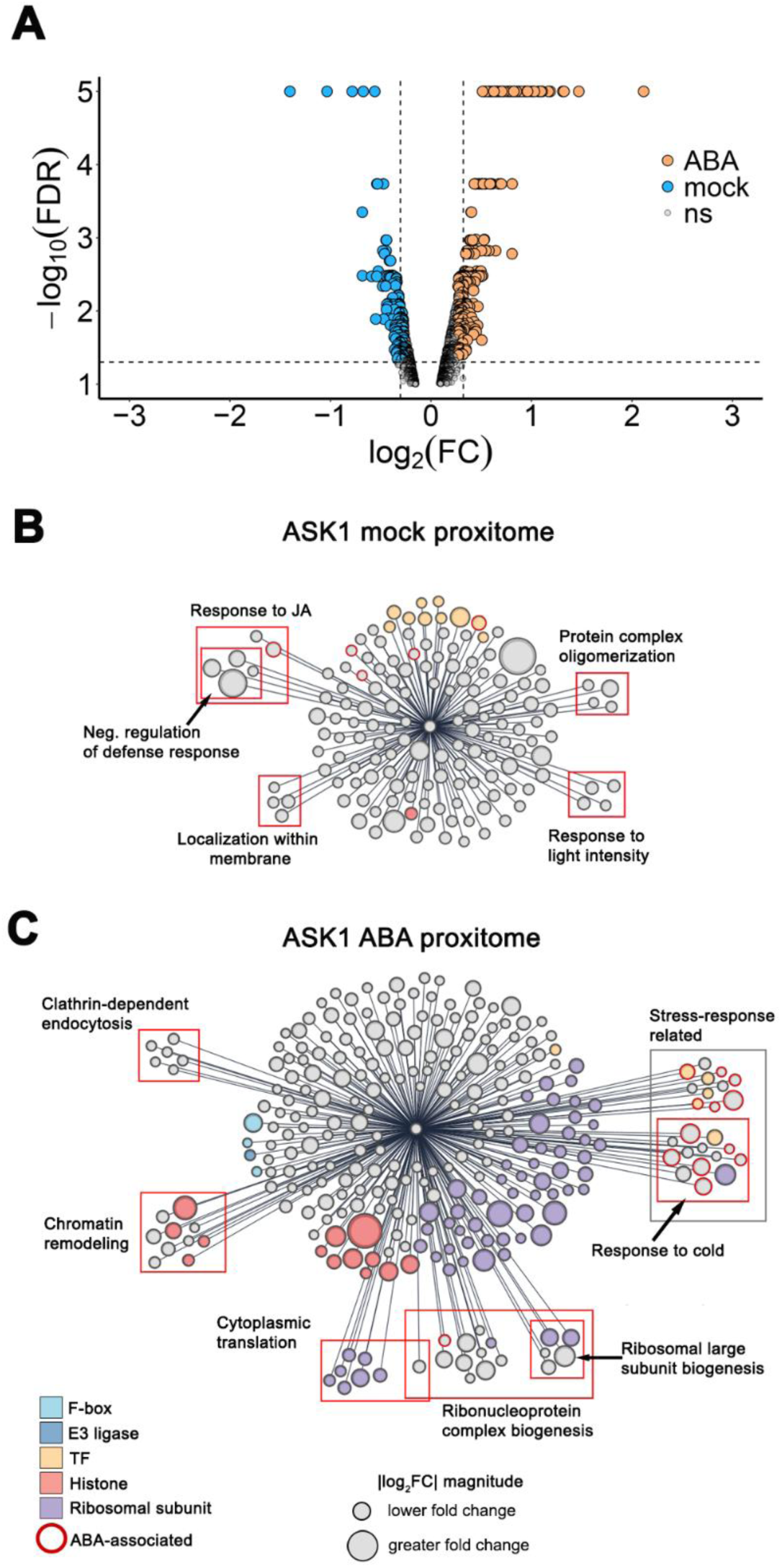
TurboID proximity labeling reveals abscisic acid (ABA) dependent remodeling of the ASK1 proxitome and enriched biological processes. **A)** Volcano plot showing proteins significantly enriched in ABA-treated TurboID-ASK1 plants compared to mock-sprayed plants of the same line (FDR < 0.05 and |log_₂_FC| ≥ 0.27, log_₂_FC values are subject to TMT ratio compression). **B** and **C)** The ASK1 proxitome detected in mock-treated and ABA-treated Arabidopsis leaves. Significant overrepresentation of gene ontology (GO) biological process (BP) annotations is shown with red boxes (P.adj <0.05). Gray boxes mark proteins with biologically relevant annotations that are not in a GO-enriched category. The size of circles represents the magnitude of enrichment (|log_₂_FC|) of each protein. Selected protein categories are color-coded within each proxitome. Proteins with ABA-related GO terms are highlighted in red.

In contrast to mock conditions, the ASK1 proxitome under ABA treatment exhibited a broad and distinct functional profile, with 30 enriched GO biological processes that were reduced to 13 categories with reduced redundancy after similarity filtering. The most significantly enriched terms were associated with ribonucleoprotein biogenesis and translation, driven by strong representation of 60S and 40S ribosomal subunits (Fig. 2C). Beyond these categories, ribosomal proteins spanning 60S, 40S, 50S, and 30S collectively accounted for ∼23% of the entire ABA enriched proxitome, whereas they were largely absent under mock conditions (Fig. 2B-c). ABA treatment also markedly enriched components of clathrin-dependent endocytosis, including dynamin-related and clathrin heavy and light chain proteins that mediate vesicle formation (Fijimoto and Tsutsumi 2014). In addition to translation and vesicle trafficking, the ABA treated plants revealed significant enrichment of abiotic stress-associated proteins, specifically of cold-responsive factors that also carry annotations for ABA, osmotic, and drought stress responses. Thirteen ABA-related proteins were enriched, comprising both positive and negative regulators of signaling. Among the positive regulators were the ABRE-binding bZIP transcription factors ABF3 and ABF4 (ABSCISIC ACID RESPONSIVE ELEMENTS-BINDING FACTORS 3 and 4 (Yoshida et al. 2010), as well as the histone deacetylases HD2A and HD2C, which promote ABA responses by repressing PP2C expression through H3K9 deacetylation (Tahir et al. 2022). Chromatin remodeling-associated proteins were strongly enriched in the ASK1 proxitome under ABA treatment, notably including multiple histones. In total, thirteen histone proteins were identified, several displaying among the highest |log_₂_FC| values relative to mock, where histones were largely absent (Fig. 2B), suggesting recruitment of chromatin-linked regulatory modules during early ABA signaling. Lastly, F-box proteins and other E3 ligases were not enriched in the mock-treated ASK1 proxitome but enriched under ABA treatment. Among these were three F-box proteins: KMD4 (KISS ME DEADLY 4), a regulator of cytokinin signaling (Kim et al. 2013); TLP2 (TUBBY-LIKE PROTEIN 2), which modulates ABA sensitivity (Jain et al. 2023); and SKIP16 (SKP1 INTERACTING PROTEIN 16), indicating that ASK1 assembles distinct SCF-type ubiquitin ligase modules in response to ABA. Together, these findings demonstrate that ABA rapidly remodels ASK1-associated complexes, recruiting chromatin regulators, translational and vesicle-trafficking factors, and a discrete set of F-box subunits that define stress-specific E3 ligase assemblies.

### Prolonged drought induces extensive remodeling of the ASK1 proxitome

To determine how ASK1 complexes reorganize during long-term water deficit, we profiled biotinylated proteins from drought-stressed and well-watered TurboID-ASK1 plants using the same TMT-based workflow applied for ABA (Fig. 1B). Across conditions, 979 ASK1-associated proteins were identified, of which 650 showed significant enrichment changes (FDR < 0.05 and |log_₂_FC| ≥ 0.27). Among the differentially enriched proteins, 338 accumulated under well-watered conditions and 312 under drought (Fig. 3A). Under well-watered conditions, functional enrichment of GO biological processes revealed ASK1 proximity to modules associated with cytoplasmic translation, vesicle organization, biotic and abiotic stress responses, and metabolism (Fig. 3B). Translation-related categories were dominated by 60S and 40S ribosomal subunits, and vesicle-trafficking factors included Golgi-tethering proteins, ESCRT-III components, and the syntaxins SYP122 and SYP132. Stress-associated terms included EXO70B2 and VPS34, which direct secretion and autophagic turnover during immune signaling (Brillada et al. 2021; Liu et al, 2020), the SA-dampening factor CBP60A (Truman et al. 2013), and the mitochondrial ROS-detox enzyme MSD1 (Holzmeister et al. 2015). ABA-linked regulators such as the positive regulator GRP3 (Shim et al. 2021) and negative regulator SF1 (Jang et al. 2014) were present, along with the Kelch F-box proteins FDR1 (Kuroda et al. 2012) and KMD3, which destabilizes PAL1 and PAL2 (Zhang et al. 2015), consistent with ASK1’s basal association with selective SCF-type modules.

**Figure 3.**
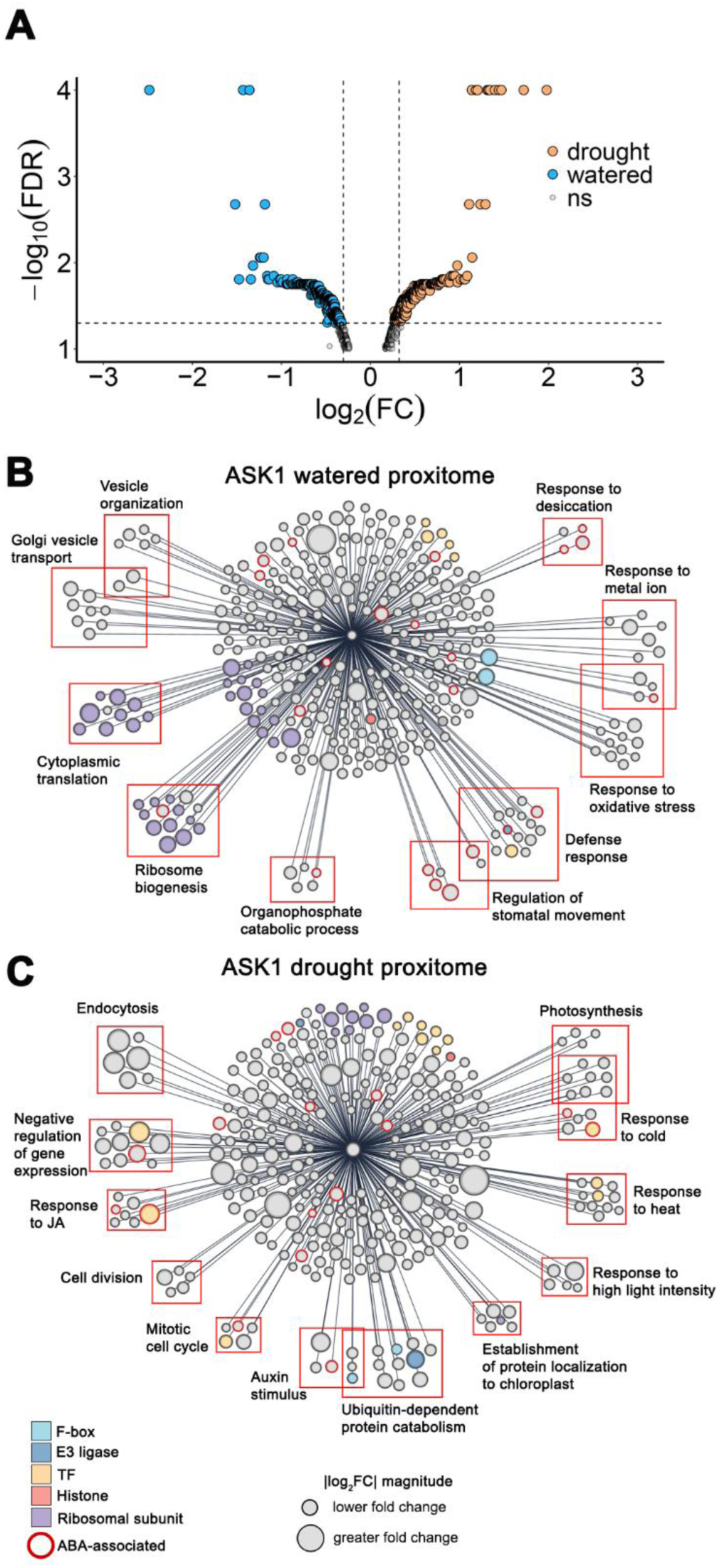
TurboID proximity labeling shows long term drought-dependent remodeling of the ASK1 proxitome. **A)** Volcano plot showing proteins significantly enriched in drought-treated TurboID-ASK1 plants compared to well-watered plants of the same line (FDR < 0.05 and |log_₂_FC| ≥ 0.27, log_₂_FC values are subject to TMT ratio compression). **B** and **C)** The ASK1 proxitome detected in well-watered and drought-treated Arabidopsis leaves. Selected significant overrepresentation of gene ontology (GO) biological process (BP) annotations is shown with red boxes (P.adj <0.05). The size of circles represents the magnitude of enrichment (|log_₂_FC|) of each protein. Selected protein categories are color-coded within each proxitome. Proteins with ABA-related GO terms are highlighted in red.

Under prolonged drought, the ASK1 proxitome shifted toward a much broader functional repertoire. GO analysis identified 156 enriched biological processes, narrowed to 45 categories with reduced redundancy after similarity filtering, compared with 24 under well-watered conditions. Drought-enriched processes spanned photosynthesis and energy metabolism, heat and light responsive signaling, protein modification and degradation, transcriptional regulation, hormone signaling, endocytosis, and cell cycle control (Fig. 3B). ASK1 was strongly associated with chloroplast import and proteostasis factors, including TOC/TIC translocon components and associated chaperones, as well as enzymes contributing to electron transport and carbon fixation. Stress-responsive modules were also enriched, prominently featuring heat- and light-responsive proteins such as the heat-shock transcription factors HSFA1B and HSFA8, which promote drought tolerance and thermotolerance (Prändl et al. 1998; Wang et al. 2020), along with their associated HSP chaperones. Among ABA linked regulators, 17 proteins carried ABA-related GO annotations, including ERD15, a negative regulator proposed to attenuate ABA signaling through posttranscriptional control of mRNAs (Kariola et al. 2006; Aalto et al. 2012), and MYC2, the most enriched positive regulator, a key bHLH transcription factor activating ABA and JA responsive genes (Abe et al. 2003; Kazan and Manners, 2013; Zeng et al. 2025).

Transcriptional repression emerged as another hallmark of the drought proxitome. The category ‘negative regulation of gene expression’ included some of the most strongly enriched drought-specific factors, notably CPL3, a C-terminal domain phosphatase-like protein implicated in modulating ABA responses through dephosphorylation of RNA polymerase II (Koiwa et al. 2002), as well as additional chromatin and RNA modifying repressors. Protein catabolism was also strongly represented: numerous 26S proteasome regulatory subunits appeared almost exclusively under drought, suggesting heightened proteasomal remodeling during chronic stress. ASK1 further recruited multiple hormone- and SCF-related components that were absent or weakly represented under well-watered conditions. These included the auxin receptor F-box protein TIR1 (Gray et al. 1999), an uncharacterized RNI-like F-box (At5g51370), and the SCF co-chaperones SGT1A and SGT1B required for TIR1-mediated substrate turnover (Gray et al. 2003). Endocytic factors also expanded strongly, most notably the dynamin-related proteins DRP2A and DRP2B, among the highest |log_₂_FC| proteins detected, which facilitate clathrin-mediated vesicle scission and contribute to cell-plate formation and polarity establishment (Huang et al. 2015). Many of these dynamin related proteins were also enriched in the ABA-dependent proxitome (Supplementary Fig. S1). Additional regulators of asymmetric cell division and cytokinesis reinforced ASK1’s association with cellular remodeling processes under drought (Fig. 3C).

Together, these findings show that prolonged drought drives extensive remodeling of ASK1-associated networks, expanding its proximity to chloroplast import machinery, proteasome regulators, transcriptional repressors, hormone-signaling components, including drought-specific F-box proteins, and endocytic and cell-cycle modules. This broad rewiring contrasts sharply with the more focused remodeling triggered by short-term ABA signaling, positioning ASK1 as a central integrator of proteostasis, signaling, and cellular adaptation during sustained water deficit.

### ASK1 overexpression alters the leaf proteome in an ABA-dependent manner

We next assessed how ASK1 overexpression influences protein abundance in response to ABA. For this, we performed quantitative protein abundance profiling using label-free Data Independent Acquisition (DIA) of WT and TurboID-ASK1 plants that were mock or ABA spray treated (Fig. 1B). TurboID-ASK1 plants overexpress ASK1 under a high expression promoter. We quantified 8,700 proteins across all samples, of which 515 were significantly different between TurboID-ASK1 and wild type under mock conditions (Q < 0.05 and |log_₂_FC| ≥ 0.27) (Fig. 4A). Among these, 205 proteins increased and 310 decreased in abundance in TurboID-ASK1 plants (Fig. 4C). Under ABA treatment, 355 proteins showed significant differences between genotypes (Fig. 4B). In contrast to mock conditions, TurboID-ASK1 plants showed more accumulation than depletion (255 accumulated; 100 depleted; Fig. 4C). Overlap between mock and ABA responsive sets was limited (12 shared accumulators; 8 shared depletions; Fig. 4D), suggesting a substantial reprogramming of the proteome in response to ABA treatment.

**Figure 4.**
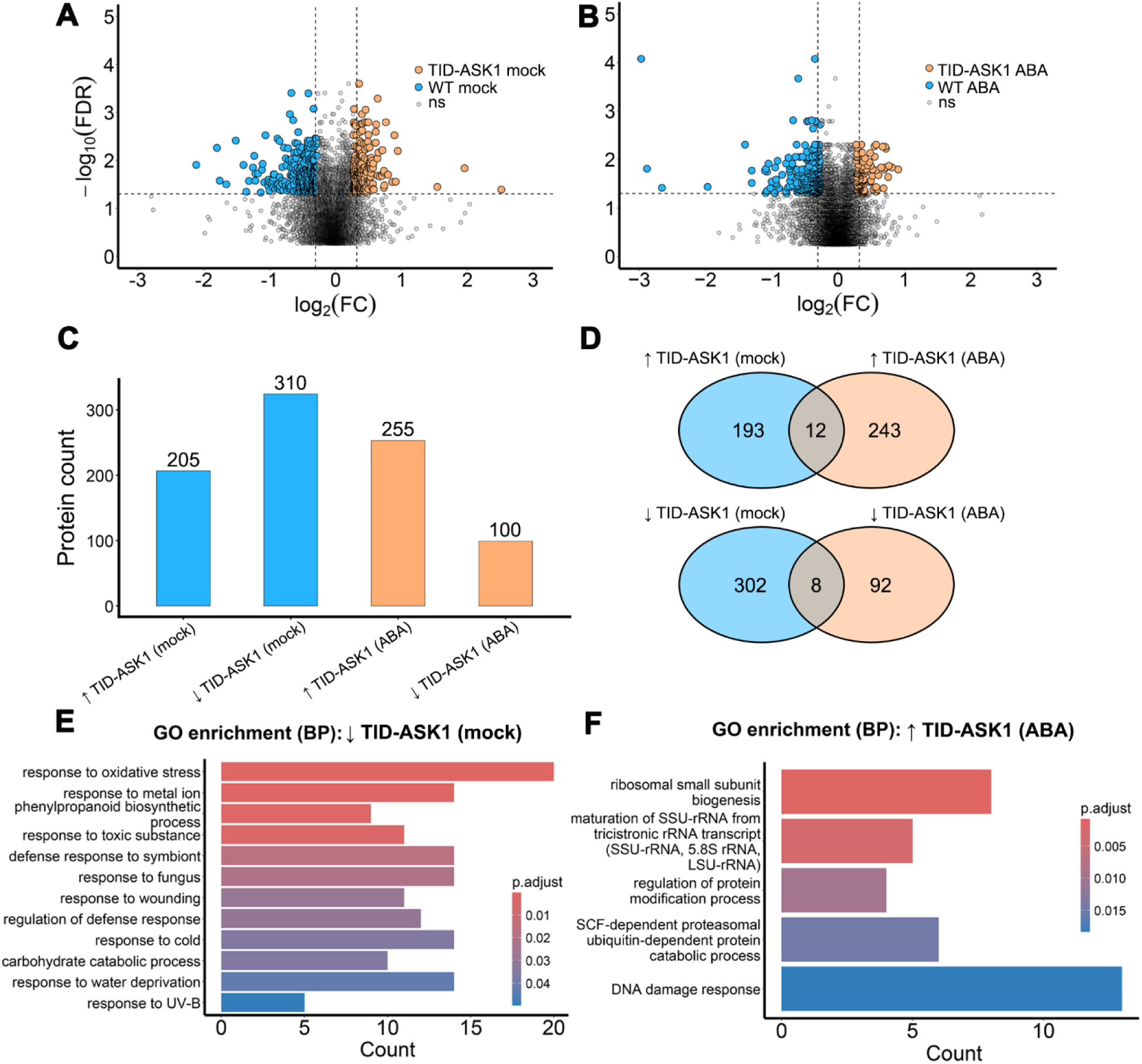
ASK1 overexpression alters the Arabidopsis shoot global proteome in an abscisic acid (ABA) dependent manner. Global proteomic analysis of ASK1-overexpressing (TID-ASK1) Arabidopsis shoots under short-term ABA spray or mock treatment, compared to wildtype (WT, Col-0). **A** and **B)** Volcano plot of proteins detected in TID-ASK1 and WT plants under mock treatment and proteins detected in TID-ASK1 and WT plants following ABA treatment. Proteins with FDR < 0.05 and absolute value of log_₂_FC ≥ 0.27 (20% change) were considered differentially abundant. **C)** Bar plot showing the number of differentially abundant proteins in TID-ASK1 compared to WT under each condition. Groups correspond to significantly increased (↑TID-ASK1) or decreased (↓TID-ASK1) proteins in TID-ASK1 plants under mock or ABA treatment. **D)** Overlaps between ↑ TID-ASK1 proteins under mock and ABA treatment (top) and ↓ TID-ASK1 proteins between mock and ABA treatment (bottom). **E** and **F)** Significantly enriched biological process GO terms in ↓ TID-ASK1 proteins under mock and ↑ TID-ASK1 proteins under ABA treatment (adjusted P < 0.05).

GO analysis revealed little functional enrichment among proteins accumulating in TurboID-ASK1 plants under mock treatment. In contrast, proteins depleted under mock were enriched in 35 GO biological processes, reduced to 12 following redundancy filtering (Fig. 4E). The majority of enriched GO terms were associated with abiotic stress and defense responses, with response to oxidative stress as the most significantly enriched. Proteins with this annotation included factors involved in antioxidant biosynthesis and several ROS-scavenging enzymes, notably the Class III peroxidases PRX37/52/53, the superoxide dismutases SOD1/2, and PER70 (PEROXIDASE 70), which showed the strongest depletion (|log_₂_FC| = 3.2). Several of these proteins also function in defense responses. Other highly depleted defense-related proteins include PR1 and PR5 (PATHOGENESIS RELATED 1 and 5), markers for SA (salicylic acid) systemic acquired resistance (SAR) (Bertini et al. 2003).

Under ABA treatment, the 273 accumulated proteins were enriched in RNA and translation pathways, together with protein modification, protein turnover, and DNA damage responses (Fig. 4F). Representative translation-related factors included DEAD-box RNA helicases such as RH27 and multiple 40S ribosomal subunits. Highly enriched DNA repair-related proteins included CHR24 (CHROMATIN REMODELING 24), NRP1 (NAP1-RELATED PROTEIN 1), and the ubiquitin E2 variants UEV1A and UEV1D involved in K63-linked ubiquitination (Xu et al. 2020). Overall, ASK1 overexpression reduces the abundance of many abiotic and biotic stress-related proteins under mock conditions, but under ABA it instead promotes the accumulation of proteins involved in translation and DNA damage repair.

### ASK1 overexpression alters the leaf proteome under prolonged drought

We next examined how ASK1 overexpression affects the global leaf proteome during long-term drought. Using DIA mass spectrometry, we quantified protein abundance in TurboID-ASK1 and wild-type plants subjected to drought (20% of soil field capacity) or well-watered conditions (Fig. 1B). Across all samples, 8,457 proteins were quantified, of which 476 showed significant differences between lines under well-watered conditions (Fig. 5A). Among these, 225 proteins were more abundant in TurboID-ASK1 plants and 251 were less abundant (Fig. 5C). Under drought there were 878 differentially abundant proteins (Fig. 5B). TurboID-ASK1 plants showed 589 depleted proteins in total, nearly twice the number that accumulated (289) (Fig. 5C). A modest overlap between proteins in drought and well-watered conditions was observed, with 31 shared accumulators and 82 shared depletions (Fig. 5D), pointing to extensive proteome reprogramming under drought.

**Figure 5.**
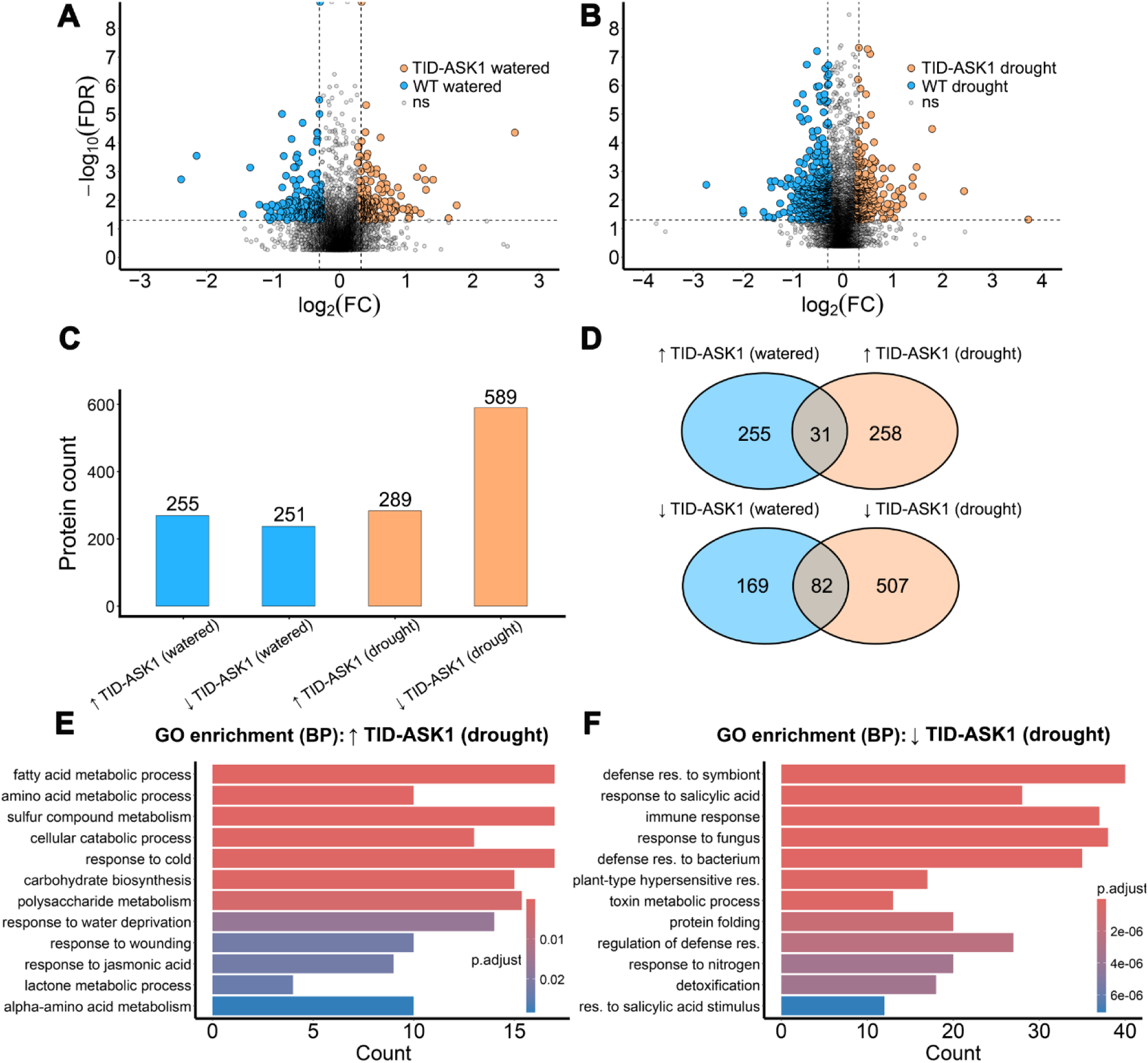
ASK1 overexpression alters the Arabidopsis shoot global proteome in a drought-dependent manner. Global proteomic analysis of ASK1-overexpressing (TID-ASK1) Arabidopsis shoots under long-term drought or well-watered conditions, compared to wildtype (WT, Col-0). **A** and **B)** Volcano plot of proteins detected in TID-ASK1 and WT plants under well-watered conditions and proteins detected in TID-ASK1 and WT plants after prolonged drought. Proteins with FDR < 0.05 and absolute value of log_₂_FC ≥ 0.27 (20% change) were considered differentially abundant. **C)** Bar plot showing the number of differentially abundant proteins in TID-ASK1 compared to WT under each condition. Groups correspond to significantly increased (↑TID-ASK1) or decreased (↓TID-ASK1) proteins in TID-ASK1 plants under well-watered or drought conditions. **D)** Overlaps between ↑TID-ASK1 proteins under well-watered and drought conditions (top) and ↓TID-ASK1 proteins between well-watered and drought conditions (bottom). **E** and **F)** Top hits for significantly enriched biological process GO terms in ↑TID-ASK1 proteins under drought and ↓TID-ASK1 proteins under drought (adjusted P < 0.05).

GO analysis identified limited functional enrichment among proteins altered in TurboID-ASK1 plants under well-watered conditions. In contrast, proteins that accumulated in TurboID-ASK1 plants under drought were enriched in 50 GO biological processes, reduced to 17 following redundancy filtering (Fig. 5E). Drought-treated TurboID-ASK1 plants showed higher abundance of proteins associated with cold, water deprivation, and defense responses, many encoded by ABA-responsive genes. Some of the most highly accumulated proteins included the stress-responsive markers RD22, RD29A, and RD29B (RESPONSIVE TO DESICCATION 22, 29A, and 29B), which are induced by both cold and drought (Yamaguchi et al. 1993; Liu et al. 2023). Proteins depleted in TurboID-ASK1 plants under drought were associated with 130 enriched GO biological processes, narrowed down to 37 after redundancy filtering (Fig. 5F). These processes were dominated by immunity and defense related functions. Reduced proteins spanned multiple tiers of the immune system, including cell surface pattern recognition, intermediate signaling components, and intracellular effector-triggered immunity pathways. Several of the most depleted proteins, however, were components of salicylic acid (SA)-mediated systemic acquired resistance (SAR). These included NPR1 (NONEXPRESSOR OF PATHOGENESIS-RELATED GENES 1), a key SAR regulator, along with its targets PR1 and PR2 (PATHOGENESIS-RELATED PROTEINS 1 and 2) (Kinkema et al. 2000). Other key proteins with reduced abundance included EDS1 (ENHANCED DISEASE SUSCEPTIBILITY 1) and PAD4 (PHYTOALEXIN DEFICIENT 4), which modulate ICS1 activity, a central enzyme in SA biosynthesis (Cui et al. 2017). SA biosynthetic enzymes themselves did not show altered abundance. Negative regulators of SA signaling were also reduced in TurboID-ASK1 plants under drought, including DMR6 and DLO1 (SA hydroxylases) indicating broad depletion across multiple components of the SA regulatory network. Overall these findings indicate that ASK1 overexpression alters canonical SCF-linked immune and SA-associated pathways as well as broader, noncanonical effects on ABA-responsive, metabolic, and stress-adaptive proteins, consistent with extensive ASK1-dependent remodeling of cellular programs under prolonged drought. Together, these complementary proximity and proteome datasets establish a coherent view of ASK1 remodeling across ABA and drought conditions, providing a basis for interpreting how ASK1 contributes to stress-responsive regulation.

## Discussion

Plants rapidly remodel signaling and proteostasis pathways to adapt to fluctuating environmental conditions, but the network-level mechanisms that coordinate these multilayered responses remain incompletely understood. ASK1, a core SKP1-like component of SCF ubiquitin ligases, has long been recognized for its canonical role in facilitating F-box-mediated substrate recognition. Classic protein-protein interaction approaches have identified many regulators of abiotic stress signaling, but these approaches are poorly suited to capture the dynamic, transient assemblies that operate in vivo during stress perception and adaptation. Here, we employed TurboID-based proximity profiling and quantitative global proteomics to follow how a central SCF adaptor reshapes its interaction landscape and associated proteome under acute ABA signaling and prolonged drought.

By validating ABA and drought responses at the physiological and transcriptional levels and then performing matched ASK1 proxitome and shoot proteome profiling under the same conditions, we obtained a coherent view of how ASK1 engages distinct molecular modules in response to short-term hormone signaling versus chronic water limitation. To our knowledge, such direct, condition-resolved mapping of a plant stress signaling hub using TurboID in intact tissues, together with whole-proteome readouts, goes beyond prior applications of proximity labeling that have largely focused on cell-type-specific or pathway-centric complexes outside defined abiotic stress regimes. Our data suggest that ASK1 may have functions extending beyond its established role as an E3 ligase adaptor. The datasets reveal that ASK1 forms highly stress-specific protein networks, recruiting distinct sets of signaling, chromatin, and proteostasis factors while displaying extensive remodeling of ASK1-dependent protein abundance in the shoot proteome.

The global proteome of ASK1 overexpressing plants shows a dramatic shift from widespread protein depletion under mock conditions to broad protein accumulation under ABA treatment, with very few proteins reduced in abundance during ABA exposure. This pattern suggests that during early stress, ASK1 may be less engaged in protein degradation and instead may contribute to the accumulation of factors involved in downstream stress responses. Consistent with this view, the ASK1 proxitome under ABA treatment showed strong associations with proteins that are not known targets SCF-mediated protein degradation.

Ribosomal subunits constituted a substantial fraction (23%) of the ASK1 proxitome under ABA treatment but were not enriched under mock conditions. In animals, ribosome ubiquitination is involved in multiple stress responsive functions, including collision sensing, nascent-chain degradation, and turnover of defective ribosomal subunits (Wooters et al. 2025). Under stress, cells downregulate ribosome biogenesis by limiting ribosomal protein synthesis and rRNA production, and increasing evidence shows that preexisting ribosomes can also be selectively degraded. For instance, in human cells subjected to starvation, the E3 ligase RNF10 ubiquitinates 40S subunits to promote their degradation, thereby adjusting ribosome abundance to reduced translational demand (Huang et al. 2025). Studies in plants are more limited, but large-scale ubiquitinome analyses indicate that ribosomal subunits are frequent ubiquitination targets (Kim et al. 2013), and several datasets support a stress-dependent component. In sugar beet exposed to salt stress, multiple 40S and 60S subunits exhibit increased ubiquitination (Liu et al. 2022), while in rice, PAMP elicitors reduce ubiquitination of ribosome associated proteins (Chen et al. 2018), suggesting that ribosome ubiquitination in plants is dynamically regulated by environmental cues. Ribosomal turnover in plants is primarily mediated through vacuolar autophagy, with no evidence supporting a role for the 26S proteasome in bulk ribosome degradation (Floyd et al. 2016; Kazibwe et al. 2019). ABA provides a relevant physiological context for this regulation, promoting broad translational reprogramming including reduced translation efficiency of ribosome biogenesis transcripts (Zhang et al. 2025) and suppression of 80S assembly with a sharp decline in polysomes (Guo et al. 2010). These effects highlight that ribosome abundance and turnover are actively modulated during stress responses, raising the possibility that SCF activity could be involved these pathways. Consistent with this idea, through our global proteome analysis we found that ASK1-overexpressing plants exposed to ABA showed elevated levels of several 40S ribosomal subunits, a pattern that has been associated with stalled translation and disrupted 80S assembly during stress (Wu et al. 2025).

However, none of the ribosomal subunits identified in the ASK1 ABA proxitome were differentially abundant in ASK1 overexpressing plants under ABA in the global proteome. In addition, a large-scale Arabidopsis ubiquitination dataset indicates that several of the ASK1-associated ribosomal proteins identified in our proxitome carry lysine ubiquitination sites (Song et al. 2024). These observations suggest that these ribosomal proteins are unlikely to be direct SCF targets for degradation under these conditions and instead support a non-canonical role for ASK1 in ribosome-associated stress responses. While it remains unknown whether ASK1 directly influences ribosome stability, the existence of stress-dependent ribosome remodeling in plants leaves open the potential for ASK1 to contribute to this process through mechanisms other than canonical proteasomal degradation. Likewise, the ABA-dependent ASK1 proxitome contained numerous strongly enriched histones, while histones were only weakly represented in the mock proxitome. Evidence from yeast and animal systems shows that histone degradation can be mediated by ubiquitination, with several E3 ligases implicated in this process. In yeast, the HECT type ligase Tom1 and the RING type ligases Pep5, Snt2, Hel1, and Hel2 have all been linked to ubiquitin-dependent turnover of core histones (Liu et al. 2005; Singh et al. 2012). In animals, histone degradation can similarly be triggered by ubiquitination via E3 ligases such as RNF8 (Xia et al. 2017) and APC/C WDR5 (Oh et al. 2020). Notably, however, no SCF E3 ligases have been reported to participate in histone turnover in any examined system. In plants, there are no known E3 ligases associated with histone degradation, and plant SCF complexes have not been linked to histone ubiquitination. In contrast, plants possess multiple E3 ligases that modify histones without degrading them. In cotton and Arabidopsis, the RING E3 ligases HUB1 and HUB2 catalyze H2B monoubiquitination, a modification that promotes chromatin relaxation and gene activation (Zhang et al. 2021; Chen et al. 2019). ABA in particular has been shown to induce several posttranslational histone modifications, including methylation, acetylation, and ubiquitination which contribute to ABA responsive transcriptional regulation (Liu et al. 2022). For example, the PRC1 components RING1A and RING1B, ubiquitinate chromatin at the ABA INSENSITIVE 4 (ABI4) locus, leading to reduced ABI4 expression (Zhu et al. 2020). However, there are no reports of SCF complexes monoubiquitinating histones in plants, animals, or yeast. Notably, the histones identified in our ASK1 ABA proxitome were not differentially abundant in ASK1 overexpressing plants under ABA treatment. Given this context, our finding that ASK1, a core SCF component, associates with numerous histone proteins under ABA treatment raises the possibility that ASK1 may engage in a non-canonical role during early stress responses, distinct from its classical function in ubiquitin-mediated protein degradation.

The global proteome of ASK1 overexpressing plants shifted from a roughly balanced number of accumulated and depleted proteins under well-watered conditions to a pronounced depletion of proteins under drought. This pattern suggests that ASK1 is predominantly associated with canonical SCF-mediated degradation during prolonged drought, with a comparatively weaker degradation role under mesic conditions. Consistent with this interpretation, the ASK1 proxitome under drought is the only condition strongly enriched for 26S proteasome subunits, as well as a larger set of highly enriched transcription factors, which are common targets of SCF and UPS mediated turnover. At the same time, however, several of the most strongly enriched proteins in the proxitome are not known targets of SCF or UPS-mediated turnover, including dynamin and endocytosis-related proteins, highlighting a potentially more complex regulatory landscape than degradation alone.

In plants, a major function of SCF complexes is to regulate transcription factors, especially those involved in hormone signaling and stress responses. The ASK1 drought-dependent proxitome showed an expanded range of transcription factors compared to the ASK1 proxitome under well-watered conditions. MYC2, also known as JA1 (JASMONATE INSENSITIVE 1), was among the most enriched transcription factors detected. MYC2 is a master bHLH transcription factor best known for activating JA dependent gene expression, but it also functions in ABA, light, circadian, and broader stress-response pathways, making it a central integrator of multiple signaling networks (Zeng et al. 2025). The SCF^COI1 complex regulates MYC2 activity indirectly by mediating the degradation of its JAZ repressors, consequently activating JA responsive transcription (Devoto et al. 2002). In contrast, MYC2 turnover is controlled directly by other ubiquitin ligases, including the U-box E3 PUB10 (Jung et al. 2015) and the CUL3-BPM E3 ubiquitin ligase complex (Chico et al. 2020), with no SCF complex known to directly target MYC2 for degradation.

Several of the most enriched proteins in the ASK1 drought-dependent proxitome are components of the endocytic machinery, a subset of which also recurs in the ASK1 ABA-specific proxitome (Supplementary Fig. S1). These include dynamin-related proteins (DRPs) that drive membrane scission through GTPase activity and adaptor proteins that recruit cargo and clathrin to form budding vesicles. The plant-specific DRP1A and DRP2B (DYNAMIN RELATED PROTEIN 1A and 2B), both enriched in the ABA and drought ASK1 proxitomes, physically interact and together support cytokinesis and cell expansion by driving constitutive endocytosis, and are involved proper PIN auxin transporter polarity (Ekanayake et al. 2021). The expression of various DRP genes is affected by cold and drought stress in plants (Duan et al. 2024), suggesting that dynamin-dependent vesicle formation may be broadly responsive to environmental stress. Moreover, large-scale ubiquitinome analyses indicate that numerous clathrin-dependent endocytic components can be ubiquitinated in planta (Berger et al. 2022), although the specific E3 ligases responsible for modifying or degrading AP-2 subunits, clathrin, DRPs, or other coat-associated factors have not been identified. The regulatory consequences of DRP ubiquitination are far better understood in animals than in plants, where dynamin-related proteins are well established to undergo ubiquitin-mediated modification and degradation. In human cells, the mitochondrial RING E3 ligase MARCH5 ubiquitinates DRP1 to promote its recruitment to mitochondria, and Parkin also ubiquitinates DRP1 to drive its proteasomal degradation. Loss of either E3 ligase disrupts mitochondrial fission dynamics (Park et al. 2010; Wang et al. 2011). DRP1 function is stress-responsive, with toxicity induced stress activating DRP1-mediated fission and cells modulating DRP1 through ubiquitination to limit damage (Li et al. 2024).

Beyond its SCF-linked functions, ASK1 may also act through SCF-independent mechanisms. The ASK1 homolog in yeast and animals, SKP1, has recently been shown to perform noncanonical functions outside of its well established role within SCF ubiquitin ligases. A defined subpopulation of SKP1 localizes to late endosome/multivesicular bodies (LE/MVBs), where nutrient-poor conditions trigger SKP1 to interact with V-ATPase subunits and promote acidification of LE/MVB-autophagosome hybrid compartments, enhancing autophagic cargo degradation (Li et al. 2023). Although there is no direct experimental evidence for an analogous pathway in plants, our proximity labeling datasets show that ASK1 becomes closely associated with proteins linked to endosomal and vacuolar fates under stress conditions. For example, the drought-dependent ASK1 proxitome includes AP2A1, a core component of clathrin-coated vesicles destined for early endosomes (Di Rubbo et al. 2013), and EPSIN1, which mediates cargo selection and membrane curvature during endosomal and vacuolar trafficking (Song et al. 2006). In addition, DRP2A, present in both the ABA and drought-dependent proxitomes, is involved in clathrin-mediated vesicle trafficking from the trans-Golgi network to the vacuole (Jin et al. 2001). In light of this context, the presence of these proteins in the ASK1 stress proxitomes may signify an SCF-mediated degradation role, a non-degradative SCF function in which ubiquitination facilitates endocytic cargo trafficking, or a distinct SCF-independent activity of ASK1 under stress.

ASK1-overexpressing plants displayed significantly reduced abundance of numerous immune and defense related proteins during prolonged drought, spanning components of both pattern-triggered immunity (PTI) and effector-triggered immunity (ETI). Notably, proteins associated with salicylic acid (SA) signaling and response were among the most strongly and broadly depleted, including central regulators of the systemic acquired resistance (SAR) pathway and negative regulators of SA signaling. These differences raise the question of why overexpressing ASK1 under drought (relative to drought-treated wildtype) would produce such a pronounced loss of SA associated immune components. One possibility is enhanced SCF-mediated turnover of key SA signaling proteins, which could prevent accumulation of both positive and negative regulators of the pathway. EDS1 and PAD4, both significantly depleted in TurboID-ASK1 under drought, form a complex needed to promote ICS1 dependent SA accumulation and to enable SA-responsive transcription. When this complex is absent, SA levels accumulate poorly and SA-dependent genes fail to activate, resulting in a dysfunctional SA defense sector (Cui et al. 2017). The BTB/POZ-domain adaptors NPR3 and NPR4, which function within CUL3 E3 ligase complexes, are known to ubiquitinate and destabilize EDS1 (Chang et al. 2019). In contrast, no E3 ligase has been reported to directly target PAD4. This leaves open the possibility that PAD4 could be subject to SCF-mediated degradation, with potentially large downstream effects on SA dependent immunity. A second possibility involves altered endocytosis-related degradation of immune receptors, consistent with the presence of clathrin-associated and endocytic proteins in the ASK1 proxitome under drought, suggesting both degradative and non-degradative SCF roles in membrane-proximal immune regulation. A third explanation is indirect. ASK1 is a positive regulator of ABA signaling (Li et al. 2012; Yapa et al. 2020), and drought elevates ABA levels, which in turn antagonize SA biosynthesis and SA responsive transcription (Tian et al. 2024). Increased ABA pathway activity in ASK1-overexpressing plants could therefore amplify ABA-SA antagonism and contribute to the broad reduction of SA associated immune proteins.

While our proximity-based approach provides a novel view of ASK1-centered protein neighborhoods (Fig. 6), the functional significance and directness of individual associations remain to be established. Future biochemical and genetic studies will be required to determine which of these ASK1-proximal proteins represent direct binding partners, SCF substrates, or components of colocated regulatory assemblies. These insights reveal adaptor plasticity as a key feature of the plant ubiquitin system and highlight ASK1 as a central integrator of transcriptional, translational, and proteostatic control during stress adaptation.

**Figure 6.**
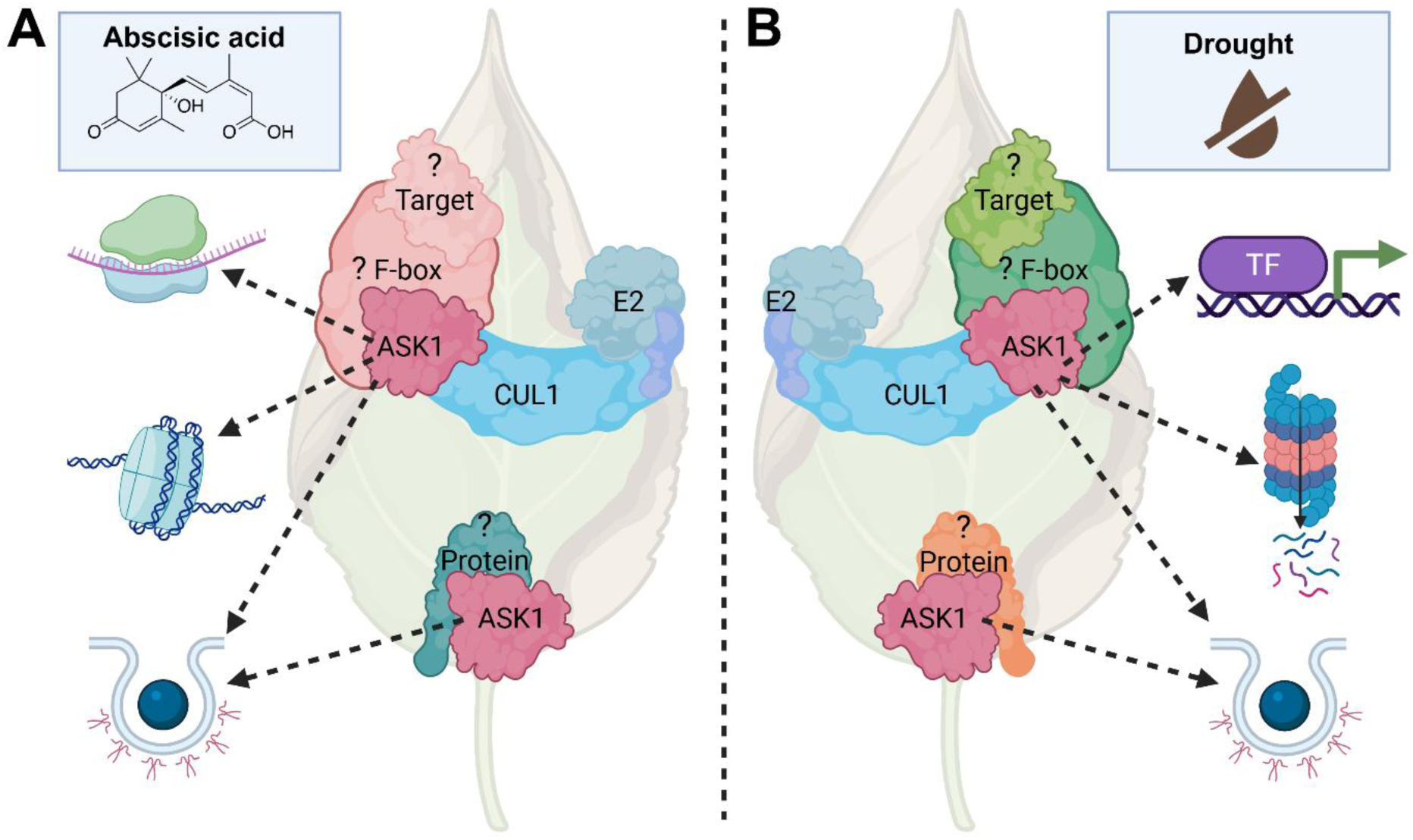
Proposed model for ASK1 as a multifunctional hub coordinating canonical and non-canonical stress responses. ASK1 serves as a plastic, multifunctional hub that integrates the plant stress responsive proteome by transitioning from rapid signaling reconfiguration under acute abscisic acid (ABA) to extensive proteostasis and cellular remodeling during prolonged drought through both canonical SCF-dependent and non-canonical pathways. **A)** Under short-term ABA exposure, ASK1 undergoes rapid proxitome remodeling, recruiting specific F-box proteins to assemble stress-specific SCF complexes. Beyond its canonical role, ASK1 establishes significant non-canonical associations with the ribosomal/translational machinery and chromatin regulatory modules, notably histones. This phase is characterized by a global shift toward protein accumulation, suggesting ASK1 may contribute to the stability of factors involved in early stress perception and downstream signaling. **B)** During sustained water deficit, the ASK1 interaction landscape expands to include a broader repertoire of transcriptional regulators (e.g. MYC2) and 26S proteasome subunits, reflecting a predominant role in canonical SCF-mediated protein degradation. In both conditions, ASK1 maintains proximity to clathrin-mediated endocytic factors and vesicle-trafficking machinery (e.g. DRP2A/B), potentially regulating the turnover or trafficking of membrane-proximal receptors and transporters. Image created with BioRender.com.

## Materials and methods

### Plant materials and growth conditions

Stable transgenic plants expressing a TurboID-ASK1 recombinant protein were previously generated in an *Arabidopsis thaliana* (Col-0) background. Plants were transformed with *Agrobacterium tumefaciens* (GV3101) containing pUBQ:TurboID-3XMyc-gASK1 through the floral-dip method, and homozygous T4 lines were selected and validated (Sun et al. 2024). TurboID-ASK1 seeds were stratified for 3 days at 4°C and transferred to Sunshine Mix #1 soil (SunGro Horticulture). Plants were kept at 22 °C, 140 µmol m^-2^ s^-1^ light intensity, and a 9 hour day, 15 hour night cycle.

### Stress treatments and physiological measurements

Five week-old TurboID-ASK1 and wildtype (Col-0) plants were treated with a 50 μM abscisic acid or mock spray. A separate group of 5 week-old TurboID-ASK1 and wildtype (Col-0) plants were allowed to dry out to 20% of field capacity, while well-watered control plants were kept at a minimum of 70% of field capacity. Soil moisture content was monitored through daily weighing of pots. For the ABA treatment assay, leaf stomatal conductance (Gs) was measured 3 hours after either ABA or mock spray treatment. For the drought treatment assay, Gs was measured upon reaching the 20% of field capacity target soil moisture. Gs was obtained from 4 randomly selected individual plants per treatment (ABA or drought) and corresponding control group (mock spray or well watered) using an Li600 Porometer/Fluorometer (LI-COR). Leaf tissue was then flash frozen for RNA and extractions for global proteomics.

### RNA extraction and transcript quantification

TurboID-ASK1 and wild-type (Col-0) Arabidopsis plants were subjected to ABA or drought treatment as described above. Leaves from three individual treated or control plants were flash frozen and pooled to generate three biological replicates per treatment and control group. Total RNA extraction was done on 100 mg of tissue per replicate, using an RNeasy Plant RNA Kit (Qiagen) following the manufacturer’s instructions. RNA (11µg) was reverse transcribed in 10µL reactions using an iScript cDNA Synthesis Kit (Bio-Rad) according to the manufacturer’s instructions. cDNA (2 µl, diluted to 1/10 with H2O) was used as template for quantitative real-time PCR amplification using PowerUp SYBR Green Master Mix (Applied Biosystems). Relative transcript levels were obtained using the comparative Ct method, normalized to *UBC21*. Primer sequences are listed in Supplementary Table S1.

### TurboID samples and preparation

For the TurboID assay, five week-old TurboID-ASK1 plants were treated with a 50 μM abscisic acid (dissolved in ethanol) or mock spray (ethanol). Leaves were then immediately syringe-infiltrated with a 50 μM biotin solution and incubated for 3 hours before flash-freezing. The selected biotin concentration and incubation time were based on previous TurboID-ASK1 protein biotinylation assays (Sun et al. 2024). A separate batch of five week-old TurboID-ASK1 plants was allowed to dry down to 20% of field capacity or maintained at 70% of field capacity as well-watered controls. Drought and well-watered plants were syringe-infiltrated with a 50 μM biotin solution upon reaching the drought target soil moisture range. Leaves were allowed to incubate for 1 hour before flash-freezing. Injected leaves from 10 treated or control plants were flash frozen, pooled, and ground to form a single replicate, with each treatment and control group consisting of three replicates. Total protein extraction was performed on 1 gram of tissue per replicate using RIPA lysis buffer, which contained 50 mM Tris (pH 7.5), 500 mM NaCl, 1 mM EDTA, 1% NP40 (v/v), 0.1% SDS (w/v), 0.5% sodium deoxycholate, 1 mM DTT, and protease inhibitor tablets (Pierce). The samples were incubated in the lysis buffer for 30 minutes at 4°C with gentle rotation, followed by centrifugation at 16,000 g for 10 minutes. The lysate soluble fraction was then loaded into Zeba™ Spin Desalting Columns (Thermo Fisher Scientific) 7K MWCO for the removal of free biotin. Desalted protein (6 mg) was then transferred to 200 μl of equilibrated Dynabeads MyOne Streptavidin C1 (Invitrogen) and incubated for 12 hours at 4°C with gentle rotation. The beads were washed in sequence with 1.7 ml each of buffer I (2% SDS), buffer II (50 mM HEPES (pH 7.5), 500 mM NaCl, 1 mM EDTA, 0.1% deoxycholic acid, 1% Triton X-100), and buffer III (10 mM Tris (pH 7.4), 250 mM LiCl, 1 mM EDTA, 0.1% deoxycholic acid, and 1% NP40), all at room temperature. Beads were then washed at 4°C, twice in 1.7 mls of 50 mM Tris (pH 7.5) and six times with 1 ml of 50 mM ammonium bicarbonate. All bead washes were performed using a DynaMag-2 magnetic rack (Invitrogen). Suspended beads (33 µL) were collected from each replicate for Western blot analysis, while the remaining sample was flash-frozen and stored at -80°C for LC-MS/MS analysis.

### Sample prep and LC-MS/MS for TurboID proximity labeling

To elute proteins from beads, samples were incubated at 95°C for 5 minutes in 1x S-Trap lysis buffer (5% SDS, 50mM TEAB, pH 8.5) with 12.5 mM biotin (Sigma; B4501). Eluted samples were digested and purified using S-Trap columns (ProtiFi; C02-micro-80), following manufacturer instructions. Samples were then reduced in 2 mM TCEP and alkylated in 50 mM iodoacetamide to disrupt disulfide bond formation. Proteins were subsequently digested into peptides at 37°C through overnight incubation with 1 µg of Trypsin and 0.1 µg of Lys-C. Peptides were then desalted using SepPak C18 columns (Waters; WAT054960). Peptides were then prepared for Tandem Mass Tag (TMTpro, Thermo Scientific) labeling through vacuum drying (Vacufuge Plus; Eppendorf) and resuspension in 0.2 M HEPES, pH 8.5 (Song et al. 2020). Hydroxylamine (1.6 µL of 5% solution) was then used to stop the TMTpro labeling reaction for each sample. Samples were incubated for 30 minutes and then pooled into one multiplex before vacuum-dying. Pooled samples were fractionated by hydrophobicity using a Pierce High pH Reversed-Phase Peptide Fractionation Kit (Thermo Scientific), with eight fractions collected. The samples were vacuum-dried and subsequently resuspended with 0.1% Optima-grade formic acid (Fisher, A117-50) in Optima-grade water (Fisher, W64). The fractions were pooled as follows: fraction 1 with fraction 5, fraction 2 with fraction 6, fraction 3 with fraction 7, and fraction 4 with fraction 8. For LC-MS/MS analysis, a 12 µL injection volume was used, containing 1.0 µg from each pooled fraction.

Chromatographic analysis was carried out using a Thermo UltiMate 3000 UHPLC RSLCnano system. A PepMap100 trap column (300 µM i.d. x 5 mm, 5 µm C18, 100 Å µ-Precolumn, Thermo Scientific) was used to desalt and concentrate peptides at a flow rate of 10 µL min-1. Sample separation was done on a 200 cm Micro-Pillar Array Column (µPAC, Pharmafluidics) with a flow rate of approximately 300 nL/min, using a 150-minute reversed-phase gradient. The gradient consisted of 80% ACN in 0.1% FA, starting from 1% to 15% over 5 minutes, 15% to 20.8% over 20 minutes, 20.8% to 43.8% over 80 minutes, then ramping from 43.8% to 99.0% over 11 minutes, and maintaining 99.0% for an additional 5 minutes. Eluted peptides were analyzed on a Thermo Scientific Q-Exactive Plus high-resolution quadrupole-Orbitrap mass spectrometer, which was directly interfaced with the UHPLC system. Information-dependent acquisition was carried out using Xcalibur 4.0 software in positive ion mode, with a spray voltage of 2.0 kV, a capillary temperature of 275°C, and an RF of 60. MS1 spectra were measured for the detection of whole-peptide mass. This was measured at a resolution of 70,000 at m/z 200, an automatic gain control (AGC) of 3e6 with a maximum ion time of 50 ms and a mass range of 350-1500 m/z. A fixed first mass of 110 m/z, an AGC of 2e5 with a maximum ion time of 120 ms, a normalized collision energy of 31, and an isolation window of 1.3 m/z were used. Charge exclusion was set to unassigned, +1, +5-8, and > +8. MS1 that triggered MS2 scans were dynamically excluded for 30 s.

### Proxitome data and gene ontology analyses

The initial dataset was processed using MaxQuant software (version 2.4.2.0) (Cox & Mann, 2008). Protein identification was achieved by searching spectral data via the Andromeda search engine against the *Arabidopsis thaliana* TAIR10 database (Cox et al. 2011; Berardini et al. 2015). MaxQuant was used to augment the proteome files with both reverse decoy sequences and a list of common contaminants. Fixed modification was set to carbamidomethylation of cysteine, while variable modifications included methionine oxidation and N-terminal acetylation of proteins. The experiment type was designated as ‘Reporter Ion MS2’, and ‘TMT18plex’ was specified for both lysine residues and N-termini. Correction factors specific to the TMT batch (TMT Lot No.: XA338617) were applied within the MaxQuant modifications tab. Enzymatic digestion was defined as ‘specific’ using Trypsin/P;LysC, allowing for a maximum of two missed cleavages. A false discovery rate (FDR) of less than 0.01 was enforced at both the peptide spectral match and protein identification levels, as calculated by MaxQuant employing a target-decoy approach (Elias and Gygi 2007). The ‘match between runs’ feature in MaxQuant was not employed. To identify significant putative interactors of ASK1, the TMT-NEAT R pipeline, incorporating the poissonseq R package, was utilized (Clark et al. 2021; Li et al. 2012). Proteins in the ASK1 proxitomes were defined based on a Q-value threshold of less than 0.05 and an absolute log2(fold change) greater or equal to 0.27.

Cytoscape (version 10.0.1) was used to visualize the ASK1 proxitome under the different conditions (Shannon et al. 2003). The R package clusterProfiler was used to test for significant enrichment of biological processes (BPs) among the identified putative interactors of ASK1 (Yu et al. 2012), with terms considered enriched if their P adjusted-value was less than 0.05. To reduce GO term redundancy and improve interpretability, we calculated a Jaccard similarity index for all GO term pairs based on the percentage of shared proteins (Ivchenko and Honov 1998). When two terms showed a Jaccard similarity of ≥0.7 (70% or more shared proteins), the term with the greater adjusted P value was removed. GO terms that were entirely encompassed by another term were also filtered out.

### Samples for leaf global proteomes

To obtain leaf global proteome data, five-week-old TurboID-ASK1 and wildtype (Col-0) plants were subjected to the same treatments used for the TurboID proximity-labeling assay: a 50 µM ABA or mock spray, long-term drought (water withheld until soil moisture reached ∼20% of field capacity), or well-watered conditions. Soil field capacity was monitored by regularly weighing pots. TurboID-ASK1 plants overexpress ASK1 under the pUBQ promoter (Sun et al. 2024). Whole shoots from ABA and mock treated plants were flash frozen 3 hours post-treatment, with three biological replicates per group. Whole shoots from drought treated and well-watered plants were collected and flash frozen when drought plants reached 15-25% field capacity, which occurred after approximately three weeks.

### Sample prep and LC-MS/MS for global proteomics

Collected tissue was ground for at least 15 minutes in liquid nitrogen using mortar and pestle. Total protein was extracted from 150 mg of finely ground tissue using the phenol-FASP method as described before (Song et al 2018, 2020, Montes et al, 2022). Finally, 1.5ug of cleaned, C18-desalted, and trypsin-digested peptides from each sample was used for LC-MS/MS.

Chromatography was performed on a Thermo Vanquish Neo UHPLC in “heated trap-and-elute, backward flush” mode. Peptides were desalted and concentrated on a PepMap Neo trap column (300 µM i.d. x 5 mm, 5 µm C18, 100 Å µ-Precolumn, Thermo Scientific) at a flow rate of 5 µL min -1. Sample separation was performed on a 110 cm Micro-Pillar Array Column (µ-PAC Neo, Thermo Scientific) with a flow rate of ∼300 nL min -1 over a 120 min reverse phase active gradient (80% ACN in 0.1% FA, from 7.2% to 24% over 109 min and from 24% to 44% in 11 min). Followed by a column/trap wash at 80% ACN for 10 min. Eluted peptides were analyzed using a Thermo Scientific Orbitrap Exploris 480 mass spectrometer with a FAIMS pro Duo interface installed, which was directly coupled to the UHPLC through an Easy Spray Ion source (Thermo Scientific). Data dependent acquisition was obtained using Xcalibur 4.0 software in positive ion mode with a spray voltage of 2.0 kV, a capillary temperature of 280 °C, an RF of 45, FAIMS compensation voltage of -45, and a total carrier gas flow of 4.2 l/min. Forty two DIA acquisition windows of 12m/z with 1Da overlap were measured at a resolution of 30,000, an automatic gain control (AGC) of 8e6 with auto maximum ion time, over a precursor mass range of 400-900 m/z. A scan range 145-1450 m/z and normalized collision energy of 29 were used.

### Proteome data and gene ontology analyses

Raw data were analyzed using Spectronaut version 19 (Biognosis). Spectra were searched, using the Pulsar search engine against the Arabidopsis thaliana TAIR10 reference annotation from The Arabidopsis Information Resource (TAIR, www.arabidopsis.org). The spectra search was performed on “direct DIA” mode. Carbamidomethyl cysteine was set as a fixed modification while methionine oxidation and protein N-terminal acetylation were set as variable modifications. Digestion parameters were set to “specific” and “Trypsin (Full)”. Up to two missed cleavages were allowed. A false discovery rate, calculated in Percolator using a target-decoy strategy, of less than 0.01 at both the peptide spectral match and protein identification level was required. Protein differential expression was assessed using the “unpaired t-test” option from Spectronaut. A Q-value of less than 0.05 and absolute value of log2(fold change) of 0.27 was used as cutoff for differential expression significance.

The R package clusterProfiler was used to test for significant enrichment of biological processes (BPs) among the significantly abundant or depleted proteins in ASK1 overexpressing plants (Yu et al. 2012). Terms were considered enriched if their P adjusted-value was less than 0.05. To reduce GO term redundancy and improve interpretability, we calculated a Jaccard similarity index for all GO term pairs based on the percentage of shared proteins (Ivchenko and Honov 1998). When two terms showed a Jaccard similarity of ≥0.7 (70% or more shared proteins), the term with the greater adjusted P value was removed. GO terms that were entirely encompassed by another term were also filtered out.

## Author contributions

D.R.-Z., J.M.-L., and N.S. conceived and designed the project. D.R.-Z., J.M.-L., N.H., C.M.-S., and S.S. performed experiments. D.R.-Z., C.M.-S., and S.S. analyzed the data. C.M.-S. and J.W. curated the data. N.S., A.T.G., and J.W. provided resources and coordinated the project. All authors contributed to manuscript writing and approved the final version.

## Data availability

Data are available via ProteomeXchange with identifier PXD072075 (ASK1-TurboID ABA), PXD072027 (ASK1-TurboID drought), PXD072105 (ASK1oe ABA), and PXDNNNNNN (ASK1oe drought)

