## Supplementary material for "Dynamic ASK1 proximity networks uncover SCF-dependent and noncanonical roles in ABA and drought adaptation": Figure S1

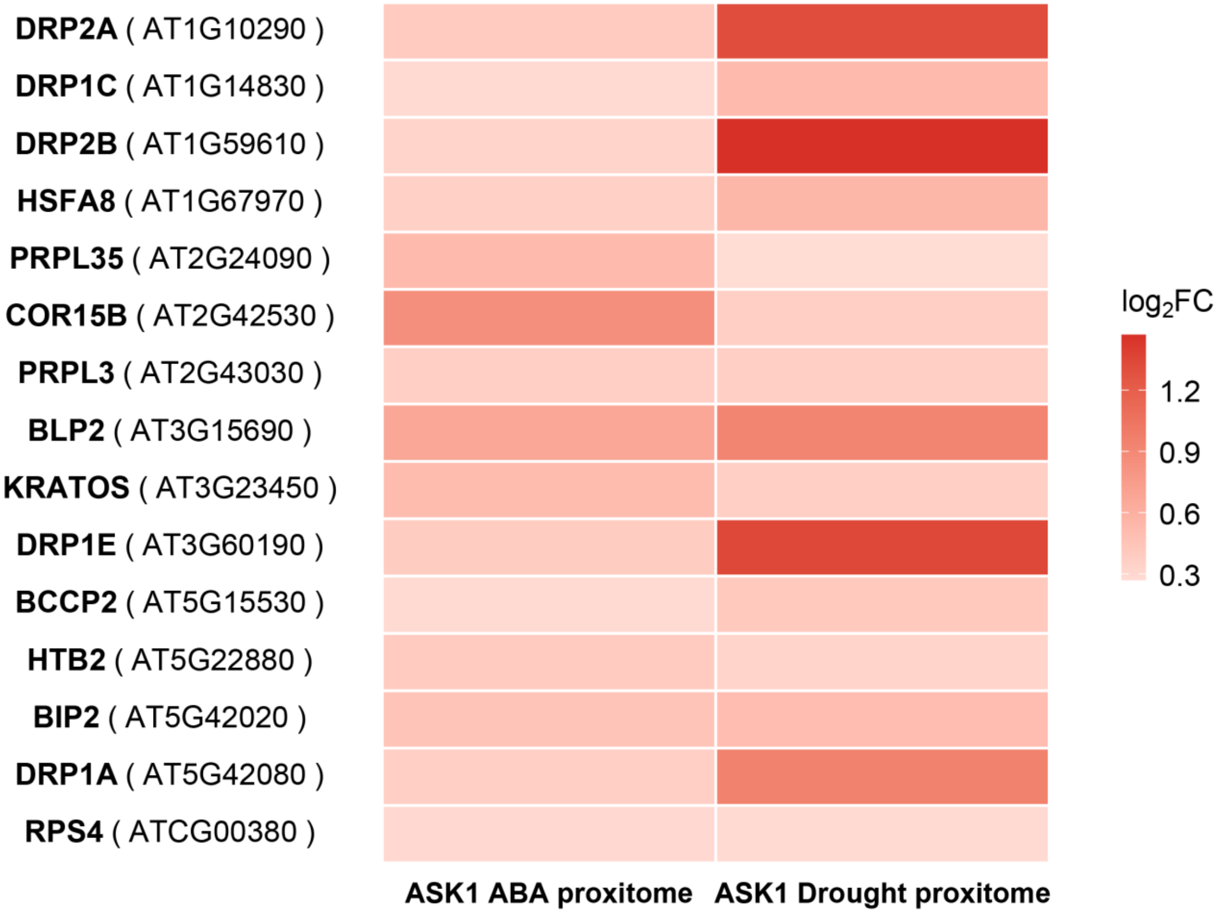


**Supplementary Figure S1.** Overlap between ASK1 ABA-dependent and drought-dependent proxitomes. Heatmap intensity indicates the magnitude of enrichment in each condition. This figure supports Figures 2 and 3.
